# Long Non-Coding RNA Transcripts in Crucian Carp Brain – Annotation and Expression Patterns in Anoxia and Reoxygenation

**DOI:** 10.1101/2025.09.07.674702

**Authors:** Magdalena Winklhofer, Göran E. Nilsson, Sjannie Lefevre

**Affiliations:** Section for Physiology and Cell Biology, Department of Biosciences, University of Oslo, Oslo, Norway

**Keywords:** Gene expression, next-generation RNA sequencing, fish, hypoxia

## Abstract

Long non-coding RNA transcripts (lncRNAs) are important regulators of various cellular processes, including gene expression. However, whether they are involved in the regulation of anoxia-induced transcriptomic changes in crucian carp (*Carassius carassius*), a species that can survive without oxygen for months at low temperatures, has remained unexplored. The existing genome annotation includes mainly protein-coding genes, which are more straightforward to annotate than the lower-expressed lncRNAs. Therefore, this study aimed to annotate lncRNAs in the crucian carp genome, investigate their potential involvement in regulating mRNA abundance in response to anoxia and reoxygenation, and further characterize any differentially regulated lncRNAs and their potential interaction partners (nearby genes). Using next-generation RNA sequencing of brain samples from crucian carp across normoxia, anoxia, and reoxygenation conditions (n=10 per group), 145,264 transcripts were assembled with a reference-guided approach. Using the bioinformatic tools CPC2, CNCI, CPAT, and FEELnc, we identified 6,072 lncRNAs among the assembled transcripts. We observed a substantial transcriptomic response to anoxia, with a total of 56,440 differentially expressed (DE) transcripts (adjusted p-value < 0.05; | log2(fold change)| > 0.38), including 1,321 lncRNAs. The FEELnc classification module identified multiple proximal RNA transcripts that may help to predict lncRNA functions, and due to their proximity to potential interaction partners (IPs), were likely *cis*-acting. Most (60%) of the interaction sites between DElncRNAs and their targets were located between genes (intergenic), predominantly on the same strand in intergenic subtypes, and primarily nested in the genic context. The log2-fold changes in RNA abundance of DE predicted IPs exhibited a positive correlation with the log2-fold changes of their corresponding DElncRNAs. These findings indicate that lncRNAs are at least co-expressed with and may regulate transcription of specific genes in crucian carp during anoxia and reoxygenation.

## Introduction

A stressor that can swiftly become life-threatening is the lack of oxygen – an essential molecule for aerobic ATP synthesis in vertebrates – and, resultingly, only a few species possess the ability to survive extended periods without it. In humans, a lack of oxygen accompanies conditions such as strokes and heart infarctions, and return of oxygen during reperfusion can worsen tissue damage (Xie, Kittur, Li, & Hung, 2022; Qin, et al., 2022). The crucian carp (*Carassius carassius*), found in small ponds in Europe and Asia, is a remarkable fish capable of surviving several months without oxygen at low temperatures, as seasonal anoxia is caused by ice and snow covering their habitat during the winter, halting photosynthesis and oxygen diffusion from the air (Holopainen, Hyvärinen, & Piironen, 1986; Vornanen, Stecyk, & Nilsson, 2009; Holopainen, Tonn, & Paszkowski, 1997). The annual life cycle of the crucian carp is thus characterized by repeated removal and restoration of oxygen access. To survive without oxygen, crucian carp reduce metabolic activity and switch to anaerobic metabolism, supported by exceptionally large liver glycogen reserves (Johansson, Nilsson, & Törnblom, 1995; Lutz & Nilsson, 1997; Haverinen, Badr, Eskelinen, & Vornanen, 2024; Boutilier, 2001). They minimize ATP use by lowering central nervous system activity and movement, but maintain cardiac output (Vornanen, Stecyk, & Nilsson, 2009). Crucially, they produce ethanol as the main anaerobic end-product, which is released into the water over the gills to prevent lactic acidosis (Lefevre & Nilsson, 2024). This is one of the central adaptations that allow them to maintain cellular energy status under oxygen-depleted conditions. However, these annual oxygen fluctuations experienced by crucian carp require temporary and reversible cellular responses. While their physiological adaptations are well-studied, the involvement and regulatory role of epigenomic mechanisms in controlling transcriptomic changes during anoxia and reoxygenation remain unclear.

The term “epigenetics” has traditionally referred to heritable changes in gene expression, occurring without any changes in DNA sequence. It has become evident, however, that the epigenetic mechanisms involved in these heritable changes in gene expression can also be induced by environmental factors, both permanent (like tissue differentiation) and temporary (like stress responses) (Dupont, Armant, & Brenner, 2009; Holliday, 1987; Wijenayake & Storey, 2016). Epigenomics, therefore, refers to mechanisms that lead to stable changes in gene expression through means other than direct regulation of transcription by transcription factors. The main forms of epigenomic regulation are DNA methylation, histone modifications, and non-coding RNAs (Li, 2021). Traditionally, RNA has been considered merely a mediator of DNA-to-protein translation, but research has revealed that some RNA molecules play versatile regulatory roles in various cellular processes (Bridges et al., 2021). Only a fraction of RNA transcripts encode proteins (mRNA), with most RNA transcripts being non-coding (St. Laurent, Wahlestedt, & Kapranov, 2015). Inamura et al. (2017) classified non-coding RNAs (ncRNAs) into housekeeping (hk. RNA) and regulatory ncRNAs (reg. RNA). Housekeeping ncRNAs include transfer RNA (tRNA), and ribosomal RNA (rRNA). On the other hand, regulatory ncRNAs encompass micro RNA (miRNA), small nucleolar RNA (snoRNA), small interfering RNA (siRNA), small nuclear RNA (snRNA), piwi-interacting RNA (piRNA), and long non-coding RNA (lncRNA) (Park & Kim, 2023; Mattick, et al., 2023; Inamura, 2017). A prior study investigated small non-coding RNAs (15-27 nucleotides) in crucian carp and other anoxia-tolerant organisms, revealing their potential role in coordinating metabolic depression and survival during anoxia (Riggs, et al., 2018). They found species-specific response patterns of conserved small ncRNA sequences, suggesting unique mechanisms of anoxia tolerance in each lineage. Due to the recent advent of small ncRNA sequencing, many novel ncRNAs have been found, but their functions remain largely unknown (Riggs, et al., 2018). To expand our knowledge of ncRNA, this study is dedicated to identifying and exploring lncRNAs. LncRNAs are RNA transcripts over 200 nucleotides in length that do not encode proteins (St. Laurent, Wahlestedt, & Kapranov, 2015; Wucher, et al., 2017; Bai, et al., 2022; Ma, Bajic, & Zhang, 2013; Ferrer & Dimitrova, 2024). LncRNAs, often expressed at lower levels than mRNAs, demonstrate specific tissue expression and are transcribed by RNA polymerase II, capped, polyadenylated, and undergo extensive alternative splicing, producing diverse functional isoforms (Bridges, Daulagala, & Kourtidis, 2021; Kazimierczyk & Wrzesinski, 2021). LncRNAs are present in both the nucleus and cytoplasm, with nuclear lncRNAs exhibiting rapid turnover to dynamically regulate gene expression and participate in chromatin interactions and nuclear organization, while cytoplasmic lncRNAs are involved in signal transduction, translation regulation, and post-translational modifications (St. Laurent, Wahlestedt, & Kapranov, 2015; Bridges, Daulagala, & Kourtidis, 2021). Hence, LncRNAs can be crucial regulators of cellular activities; interacting with DNA, mRNA, and proteins to modulate transcription, epigenomic modifications, chromatin remodeling, splicing, translation, and post-translational modifications (Bridges, Daulagala, & Kourtidis, 2021; Kazimierczyk & Wrzesinski, 2021; Wang, et al., 2014).

Functionally, lncRNAs appear to regulate gene expression in two distinct ways: locally (*cis*) and distantly (*trans*). *Cis*-acting lncRNAs function based on their transcriptional site, and *trans*-acting lncRNAs act elsewhere, independent of their transcription site (Ma, Bajic, & Zhang, 2013; Du, Zhang, Wu, You, & Dong, 2023). *Cis*-acting lncRNAs can either enhance target gene expression by attracting activating factors or silence gene expression by inhibiting the promoter by recruiting repressive complexes (Kazimierczyk & Wrzesinski, 2021; Park & Kim, 2023). They can also inhibit transcription by overlapping with target genes, blocking complex assembly, and thereby inhibiting the initiation of expression (Song, Wang, Zhu, & Dong, 2019). They can act as natural antisense RNAs to promote mRNA degradation and regulate gene expression post-transcriptionally by binding to splicing factors or hybridizing with mRNA sequences to block translation (Song, Wang, Zhu, & Dong, 2019; Ma, Bajic, & Zhang, 2013). *Trans*-acting lncRNAs, on the other hand, bind to proteins, DNA, and RNA; interacting with RNA-binding proteins to function as posttranscriptional factors, inhibiting mRNA splicing, altering stability, and affecting translation (Bai, et al., 2022). Understanding whether lncRNAs are *cis* or *trans*-acting is essential for understanding their regulatory mechanisms and roles in gene expression. A single lncRNA can interact with multiple target genes, while multiple lncRNAs can collectively regulate a single target gene (Park & Kim, 2023). They influence development, cell cycle, differentiation, metabolism, stress response, inflammation, cancer, and apoptosis (Song, Wang, Zhu, & Dong, 2019; Ma, Bajic, & Zhang, 2013; Bai, et al., 2022; Statello, Guo, Chen, & Huarte, 2020; Barreca, Zichittella, Alessandro, & Conigliaro, 2021).

This study examined the transcriptomic changes in the brain from normoxia to anoxia and reoxygenation, aiming to identify and annotate lncRNAs in the crucian carp genome. We investigated their abundance, identified potential interaction partners, and examined the genomic context in which they appear in relation to their potential interaction partners. Furthermore, we assessed the nature of these interactions in terms of whether they are likely to be *cis*- or *trans*-acting and investigated if changes in the RNA abundance of the identified lncRNAs and the mRNA abundance of their potential interaction partner were correlated. These data enhance our understanding of the physiological and molecular responses to oxygen stress and improve the functional annotation of the crucian carp genome.

## Materials and Methods

Two distinct batches of samples were utilized in this study: batch 1 with n=6 per condition and batch 2 with n=4 per condition. Both batches were processed using identical protocols. However, the primary variation between them lay in the sequencing technique and depth employed. To account for the methodological differences, the batches were analyzed separately, but both were used to identify common lncRNAs with CPC2, CNCI, CPAT, and FEELnc. Due to the methodological differences, only batch 1 was analyzed for differential expression analysis. This batch was chosen because of its larger sequencing depth and thus coverage. Details of the exposure for the fish in batch 1 are provided below, while information on the fish in batch 2 is provided in (Winklhofer, Andersen, & Lefevre, 2025), with the exposure and sampling methods being consistent with those used for batch 1.

### Crucian Carp Husbandry and Exposure Experiment

Crucian carp (n = 6 per condition, both sexes, mean mass = 28.84 ± 1.65 *g*) involved in this study were obtained from the Tjernsrudtjernet pond (Oslo, Norway) in September and kept in a 750-liter holding tank in the aquarium facility at the Department of Biosciences, University of Oslo. The holding tanks were supplied with aerated de-chlorinated Oslo tap water at 9 - 10°C and subjected to a 12 h darkness / 12 h daylight cycle. The animals were fed daily with commercial carp food and maintained for 15 months under stable conditions to acclimatize them to the facility and evaluate their health and feeding behavior before the experiments were conducted. The anoxia-exposure experiments were conducted according to the Norwegian animal research guidelines (“Forskrift om bruk av dyr i forsøk“) at an approved animal facility (Norwegian Food and Safety Authority, approval no. 155/2008, FOTS ID 16063). To replicate natural conditions, fish were not fed during the 24-hour acclimatization and exposure period, as feeding could introduce variability and degrade water quality (Holopainen, Hyvärinen, & Piironen, 1986; Nilsson, 1990). A flow-through system filled two 25-liter buckets with tight lids with aerated water. One bucket served as the normoxic control, while the others were subjected to anoxia or reoxygenation. Each contained 25 fish, ensuring consistent conditions, except for oxygen supply, which was monitored daily with a WTW Oxi3310 oxygen meter. A total of six biological replicates were sampled for each condition for this study, while the remaining fish were allocated for other research purposes within our group. The consistent fish counts across all tanks ensured uniform conditions (biomass), except for variations in oxygen levels. Food was withheld throughout the exposure. The normoxic bucket maintained above 95% air saturation through air bubbling. Nitrogen replaced the air bubbling in the anoxia and reoxygenation buckets. The oxygen levels of the anoxic buckets were below the meter’s detection limit and thus considered anoxic (below 1% air saturation). After anoxia, the reoxygenation bucket resumed air bubbling. Fish in normoxic and anoxic conditions were sampled on the seventh day (n=6 per condition), and the reoxygenation group was sampled after an additional 24-hour reoxygenation period (n=6). During sampling, each fish was individually netted and euthanized with a sharp blow to the head, followed by cervical transection. The brain tissue was extracted within one minute by making an incision towards the eyes, severing the olfactory tract, optic nerve, and spinal cord to remove the entire brain in one piece. The brain tissue was immediately frozen in liquid nitrogen and stored at -80°C until RNA extraction and sequencing.

### Total RNA Extraction, Quality Assessment, cDNA Library Preparation, and Sequencing

Total RNA was extracted from each brain sample using TRIzol® Reagent (Ambion RNA, Life Technologies) according to the manufacturer’s protocol, without DNase treatment. The RNA concentration measured by Nanodrop was 1359.2 ± 360.0 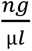 (mean ± s.d.) (batch 1) and 751.29 ± 350.63 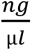 (mean ± s.d.) (batch 2). Total RNA samples were submitted to the Norwegian Sequencing Centre (NSC) at the University of Oslo’s Department of Biosciences for further quality control, library preparation, and sequencing. RNA concentration and quality were assessed using an Agilent 2100 Bioanalyzer with a Eukaryote Total RNA Nano assay. For the batch 1 samples, the total RNA concentration was 895.7 ± 227.7 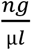 (mean ± s.d.), the rRNA Ratio [28s/18s] was 1.8 ± 0.1, and the RNA Integrity Number (RIN) was 8.5 ± 0.3 (measured in 11 of 18 samples; the rest lacked an RIN due to overloading). The RNA quality was deemed sufficient for library preparation by the NSC. The cDNA library was prepared using the Illumina Strand-specific TruSeq™ RNA-seq kit (enrichment of poly-(A)-tailed mRNA). Sequencing was performed on an Illumina HiSeq 2500, with 18 samples (six per treatment) multiplexed on four lanes, producing 125 bp paired-end reads with an expected sequencing depth of a minimum of 40 M reads per sample.

### Pre-Processing of RNAseq Reads and Transcriptome Assembly

The raw RNA data included 125 bp (batch 1) and 150 bp (batch 2) paired-end reads that underwent quality assessment using FASTQC (v0.11.8-Java-11) and adapter clipping and quality trimming utilizing TrimGalore (v0.3.3) with the settings “--quality 20 --length 40”. To align the reads to the crucian carp genome (Valencia-Pesqueira, Hoff, Tørresen, Jentoft, & Lefevre, 2025), STAR (v2.7.11b-GCC-13.2.0; (Dobin, et al., 2012)) was used with the settings “ --runThreadN 8 --twopassMode Basic --outSAMtype BAM SortedByCoordinate -- limitBAMsortRAM 3000000000 --outBAMsortingThreadN 5 --outSAMattributes All -- outSAMstrandField intronMotif”. The analysis was run separately for batch 1 and batch 2 samples, to accommodate indexes suited to the different lengths. The genome for the batch 1 samples (star_index_125) and batch 2 samples (star_index_150) were indexed using the settings “--runMode genomeGenerate --sjdbOverhang 124” and “--runMode genomeGenerate --sjdbOverhang 149” respectively, with the genome file “ccar_genome_v1_262scaffolds.fasta” and the annotation “carcar_annotation_v5.gtf” provided. To assemble the transcriptome, we used StringTie, a genome-guided transcriptome assembly tool that incorporates *de novo* genome assembly concepts to enhance transcript accuracy and completeness (Pertea M., et al., 2015). StringTie (v2.2.1-GCC-11.2.0) (Pertea M., et al., 2015) was executed with default settings, utilizing the annotation file “carcar_annotation_v5.gtf” (Valencia-Pesqueira, Hoff, Tørresen, Jentoft, & Lefevre, 2025) to generate a transcriptome for each biological sample. These individual transcriptomes were subsequently merged using the StringTie “--merge” command to create a comprehensive transcriptome that included all biological samples from both sample batches (see Supplementary Material 01).

### Annotation of LncRNAs

Transcripts from the assembled transcriptome were extracted using GFFread (v0.9.0; default settings) with the genome “ccar_genome_v1_262scaffolds_masked.fasta” provided (see Supplementary Material 02) (Pertea & Pertea, 2020). The pipeline employed was developed following the one used in Kang et al. (Kang, et al., 2024). The programs CPC2 (v1.0.1; Biopython: v1.81-foss-2022b), CPAT (v3.0.4), CNCI (v2), and FEELnc (v.0.2.1) were employed to determine the coding potential and identify lncRNAs. As recommended by the CPC authors, these programs were selected to utilize their diverse algorithms and approaches for comprehensive re-analysis (Kang, et al., 2017). Alignment-based tools compare sequences to reference genomes, excelling at identifying conserved protein-coding genes. However, they can be biased towards well-conserved sequences and may miss species-specific lncRNAs (Wang, et al., 2013). Alternatively, alignment-free tools analyze k-mer frequencies or other statistical properties, enabling the detection of unique or species-specific lncRNAs without relying on direct sequence alignment (Kang, et al., 2024). Given the challenges of non-model organisms with lineage-specific and less conserved lncRNAs, we used primarily species-neutral, alignment-free tools. Alignment-based methods can be biased towards conserved protein-coding genes and struggle to detect species-specific lncRNAs, limiting their effectiveness (Wang, et al., 2013). CPAT, released in 2013, runs on an alignment-free logistic regression model (Wang, et al., 2013). The program, trained with "zebrafish_Hexamer.tsv" and "Zebrafish_logitModel.RData," was executed using the GFFread output file containing all transcripts (“02_gffread_cc_transcripts_all_fa.txt”; see Supplementary Material 02) and the setting “–top-orf=1” to output only the best open reading frame (ORF) hits, with a non-coding transcript cut-off of ≥ 0.381 (= cutoff for long non-coding transcripts when training with zebrafish data, as stated by the program developers) (see Supplementary Material 03). Further, we employed CNCI, which profiles adjoining nucleotide triplets to distinguish protein-coding from non-coding sequences. Their program developers describe CNCI as being tailored towards incomplete transcripts and sense-antisense pairs and performing well across vertebrates, leveraging intrinsic sequence composition (Sun, et al., 2013). CNCI was provided with the GFFread output file containing all transcripts (“02_gffread_cc_transcripts_all_fa.txt”; see Supplementary Material 02) and ran with the “-m ve” model for vertebrates to identify lncRNAs (see Supplementary Material 04). Considering the advancements in lncRNA research since the release of CPAT and CNCI nearly a decade ago, we included newer tools like CPC2 and FEELnc, introduced in 2017, to reflect these developments (Kang, et al., 2017). CPC2 is a species-neutral, alignment-based tool that uses intrinsic sequence features and a hierarchical feature selection procedure with a random forest function and ten-fold cross-validation to determine coding probability (Antonov, Mazurov, Borodovsky, & Medvedeva, 2018). The developers of CPC2 state that it is well-suited for expanding transcriptomes of non-model organisms, offering balanced classification of protein-coding and non-coding transcripts (Kang, et al., 2017). We used standard settings to provide the GFFread output file containing all transcripts ("02_gffread_cc_transcripts_all_fa.txt"; see Supplementary Materials 02 and 05). FEELnc (Flexible Extraction of Long non-coding RNA) is an alignment-free tool that uses a Random Forest model to distinguish lncRNAs from mRNAs based on RNA size, ORF coverage, and k-mer usage (Muret, et al., 2017; Antonov, Mazurov, Borodovsky, & Medvedeva, 2018). Trained with nucleotide frequency patterns and relaxed ORFs, it effectively separates lncRNAs from protein-coding RNAs by capturing diverse characteristics (Schneider, Raiol, Brigido, Walter, & Stadler, 2017). FEELnc employs a shuffle strategy to model species-specific lncRNAs without a training set, making it versatile across species (Wucher, et al., 2017). Its main advantage is automatically computing an optimal cut-off for enhanced prediction sensitivity and specificity and classifying lncRNAs by genomic position and orientation relative to reference genes (Muret, et al., 2017). FEELnc consists of three modules: filtering, coding potential, and classification. We used the assembled transcriptome ("01_Stringtie_cc_transcriptome_merged_gtf.txt") with the setting "--monoex=1" to retain all monoexonic transcripts, along with the annotation "carcar_annotation_v5.gtf" (Valencia-Pesqueira, Hoff, Tørresen, Jentoft, & Lefevre, 2025) for filtering. The filtered candidates were then analyzed for coding potential using the setting "--mode=shuffle" with the same annotation “carcar_annotation_v5.gtf” and the genome "ccar_genome_v1_262scaffolds_masked.fasta" (see Supplementary Material 06) (Valencia-Pesqueira, Hoff, Tørresen, Jentoft, & Lefevre, 2025). Following the prediction of lncRNAs, only those consistently recognized by CPC2, CNCI, CPAT, and FEELnc were retained for further analysis, using Python for filtering (see Supplementary Materials 07).

### Prediction and Characterization of lncRNA Interaction Partners

The lncRNAs identified by FEElnc ("06_feelnc_cancidate_lncRNA.gtf_RF.txt"; this file holds the final set of lncRNAs, and does not include other transcripts just lacking ORFs) together with the annotation "carcar_annotation_v5.gtf" were subsequently used in the classifier module of FEELnc to predict interaction partners (see Supplementary Material 08). This module employed a sliding-window approach to identify potential overlaps with the nearest transcripts from the reference annotation. The classification process was conducted in two layers, with the first layer distinguishing between genic and intergenic regions. In contrast, the second layer identified subtypes based on interaction orientation to determine their spatial localization. Firstly, the lncRNAs (“08_lncRNA_classes.txt”; see Supplementary Material 08) were filtered to only retain the lncRNAs that were identified by all four algorithms (CNCI, CPC2, CPAT, FEELnc; see Supplementary Material 07). For each lncRNA interaction, the optimal lncRNA-to-RNA-partner pair was identified and labeled as “isBest” (by FEELnc classifier module), these pairs were subsequently filtered (with Python). The best partner for intergenic interactions was selected as the one closest to the long intergenic non-coding RNA (by the FEELnc classifier module). The best RNA partner in genic interactions was chosen based on a prioritization principle of exonic, intronic, and containing transcripts (by the FEELnc classifier module) (Wucher, et al., 2017). The classified lncRNAs (“lncRNA_classes.txt”) were further categorized into lncRNAs or DElncRNAs (see section *Differential Expression of LncRNAs below*). Subsequently, all visualizations and plots were generated using Python.

### Differential Expression of LncRNAs

We used only the BAM files from batch 1 for all the following analyses, as they provided the highest and sufficient coverage and sequencing depth. This approach eliminated the complications associated with normalizing data from different sequencing libraries with 125 bp and 150 bp reads, as well as addressing variations in sequencing depth and other potential batch effects. The aligned BAM files (aligned with STAR and transcriptome assembled with StringTie; see section *Pre-Processing of RNAseq Reads and Transcriptome Assembly above*) were utilized in featureCounts, a module of Subread (2.0.3-GCC-11.2.0), with the settings “-T 4 -O -C -p --countReadPairs -s 2 -t exon -g transcript_id” to generate exon counts summarized at the transcript levels from the RNA sequencing data (see Supplementary Material 09). This configuration allowed multi-overlapping reads (for transcripts) but discarded chimeric alignments. Differential expression analysis was conducted using DESeq2 (v1.40.2) in R (see Supplementary Material 10 – 12). Identifying differentially expressed transcripts was performed in Python by filtering out rows with fewer than ten counts, applying a | log2(fold-change) | threshold of 0.38 (corresponding to a fold-change larger than 1.3 and smaller than 0.77), and selecting those with an adjusted p-value ≤ 0.05. Of the initial 6,072 lncRNAs, only 5,016 met the threshold of ten or more read counts and were subsequently utilized for plotting (in Python).

### Differential Expression of DElncRNA Interaction Partners

The aligned BAM files (samples from batch 1; aligned with STAR to the crucian carp genome; see section *Pre-Processing of RNAseq Reads and Transcriptome Assembly above*) were utilized in featureCounts, a module of Subread (2.0.3-GCC-11.2.0), with the settings “-T 4 -C -p -- countReadPairs -s 2 -t exon -g gene_id” to generate exon counts summarized at the gene levels from the RNA sequencing data (see Supplementary Material 13). Differential expression analysis was conducted using DESeq2 (v1.40.2) in R (see Supplementary Material 14 – 16) on annotated genes after filtering out genes with fewer than ten counts. Identifying differentially expressed genes was performed in Python by applying a | log2(fold-change) | threshold of 0.38 (corresponding to a fold-change larger than 1.3 and smaller than 0.77), and selecting those with an adjusted p-value ≤ 0.05. All subsequent plotting was performed in Python. To visualize the mRNA abundance patterns of the predicted interaction partners, of DElncRNAs, which were differentially expressed themselves (DEIPs), we utilized the feature count gene matrix to extract the abundance data, omitting rows with fewer than 50 total counts across all samples. A cluster map was generated with z-score scaling to normalize row-wise expression. Row clustering was implemented to identify gene clusters with similar expression patterns, while column clustering was disabled to preserve the experimental sequence of conditions: normoxia, anoxia, and reoxygenation. To explore whether columns aligned with their experimental conditions, a PCA plot was constructed using Python. A joint plot was generated to examine the expression correlation between the DElncRNAs and their corresponding predicted interaction partner under varying experimental conditions. The log2(fold-changes) from the DESeq analysis were utilized for both the DElncRNA at the transcript level (based on the gene count matrix derived using the newly constructed transcriptome via StringTie), and their predicted interaction partners (at the gene level by using the gene count matrix from the annotation that includes only protein-coding genes). To statistically support the correlation, the Pearson correlation was calculated using Python. Furthermore, we filtered the differential expression data to visualize the top three most significant changes, characterized by the smallest adjusted p-values and largest log2 fold changes (not considering their experimental condition comparison), for DElncRNA and predicted interaction partners. To ensure reliability, candidates with fewer than 50 cumulative counts across all samples were excluded based on the feature count gene matrix counts. Subsequently, TMM-normalized abundance (counts per million) was computed (with the conorm package in Python) and normalized against the mean normoxia expression, from which relative deviations were calculated. The overall effect between the treatment groups was assessed using a one-way ANOVA when standard deviations were consistent (within 0.5 of the mean). Otherwise, the Alexander-Govern test was employed. Significant pairwise differences were identified through a Tukey post-hoc test in both scenarios. To explore the differentially expressed potential interaction partners of DElncRNAs identified by the FEELnc classifier (for more information, see *Prediction and Characterization of lncRNA Interaction Partners*) and to understand their possible coding functions or involved processes, gene ontology (GO) enrichment analysis was performed using the goseq package (v1.52.0) (see Supplementary Material 17).

## Results

### Identification and Differential Expression of Long Non-Coding Transcripts

Sequencing yielded around 40 million (batch 1) and 16.7 million (batch 2) read pairs per sample. The mapping efficiency (reads uniquely aligned and properly paired) of these clean reads using STAR was 95.45 ± 1.58% (batch 1) and 90.94 ± 3.62% (batch 2). Principal component analysis (PCA) of the mRNA abundance data (batch 1) showed clustering of the six biological replicates according to their experimental conditions (Figure 1A). No significant differences were observed between male and female samples, indicating no sex-specific variation in gene expression. Using StringTie, we identified 145,264 transcripts, with their length distribution shown in Figure 1B (green). We used the CPC2, CNCI, CPAT, and FEELnc pipelines to identify 6,072 long non-coding transcripts common to all four programs (Figure 1C), none of which were already present in the annotation of the crucian carp genome (see Supplementary Material 18). All transcripts were used to investigate differentially expressed transcripts (DETs) across three comparisons [normoxia to anoxia (N to A), normoxia to reoxygenation (N to R), and anoxia to reoxygenation (A to R)]. The 56,440 DETs exhibited a similar length distribution compared to all transcripts (Figure 1B; blue) and hundreds of DETs were identified in each comparison, with roughly equal numbers of up- and down-regulated transcripts. The highest number of transcriptional changes was observed in the N to A comparison (Figure 1D). In contrast, the A to R comparison showed the fewest changes, while the N to R comparison displayed numbers similar to the N to A comparison. Further, we examined the overlap of DElncRNA transcripts identified in the DET analysis across the three comparisons (Figure 1E). Most of the 1,321 DElncRNAs in the N to A comparison overlapped with the N to R comparison, and there was a notable overlap between the N to A and A to R comparisons. Furthermore, the smallest overlap between two conditions of DElncRNA transcripts was shared between the N to R and A to R comparisons.

**Figure 1:**
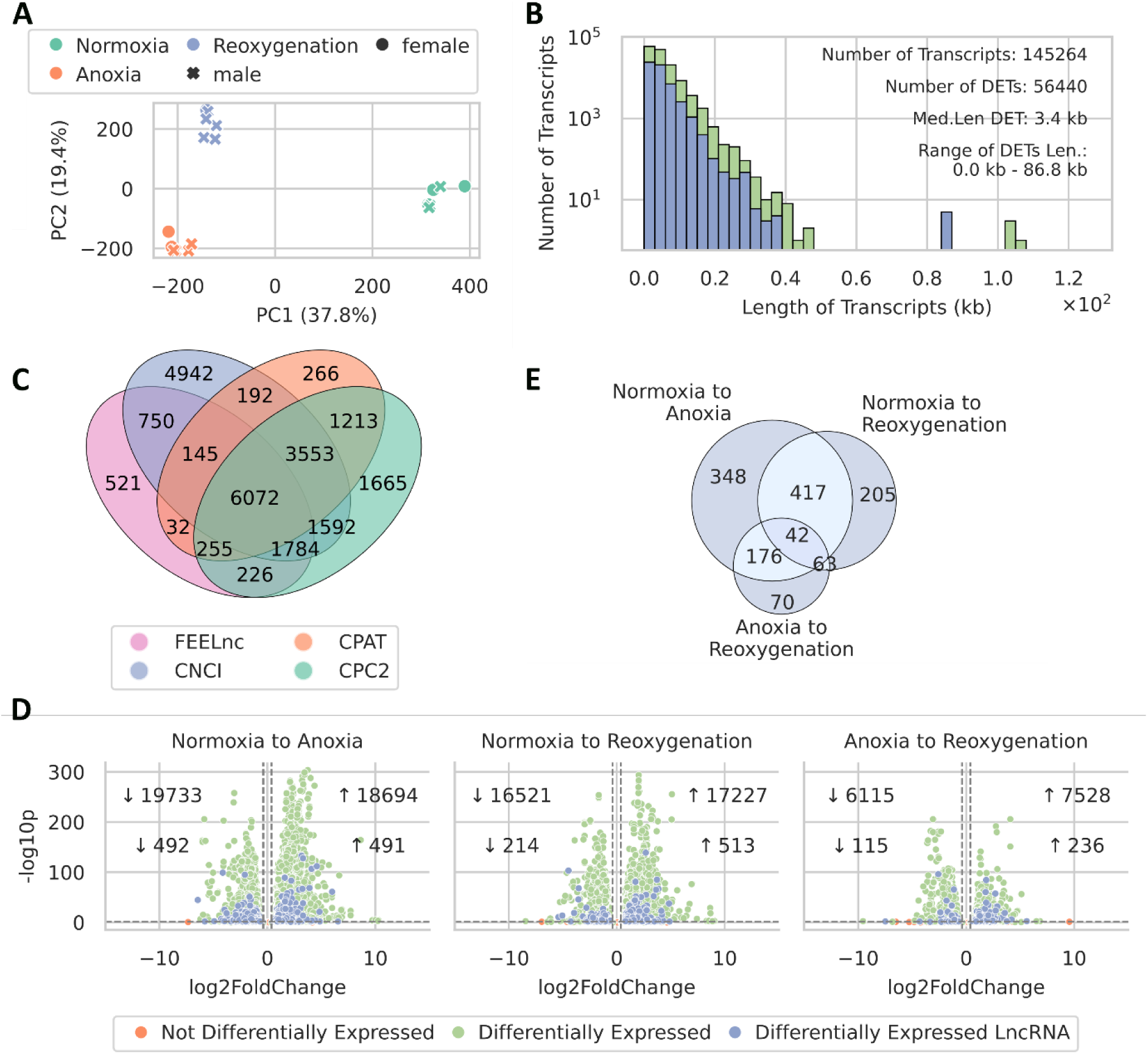
Transcriptome of the different conditions and comparisons. A: PCA of sample clustering in the RNA sequencing data in the six biological replicates for the three conditions (batch 1). B: Length distribution of all transcripts (green) and DE transcripts (blue) in the genome. The number, median length, and range of DE transcripts are given. Lengths are in kilobases (kb). C: Number of lncRNAs identified by FEELnc, CPAT, CNCI, and CPC2. Overlaps indicate lncRNAs found by multiple programs, highlighting consensus and unique findings (including samples from batch 1 and batch 2). D: Significantly DElncRNA transcripts (batch 1). The plot highlights DElncRNA transcripts between the treatment conditions, using adjusted p-value < 0.05 and a |log2(fold-change) | threshold of 0.38 (corresponding to a fold-change larger than 1.3 and smaller than 0.77). The top number in each inset indicates the number of differentially expressed transcripts (green). The bottom number indicates numbers for DElncRNAs (blue) with increased (↑) or decreased (↓) expression in each comparison. E: Venn diagram of overlapping DElncRNA transcripts between the comparisons.

To contextualize our genome annotation information, we compared the number of genes and transcripts and identified lncRNAs, as well as their median, minimum, and maximum lengths, to other fish genomes in the Cyprinidae family. Detailed listings of extracted genome annotations, reference numbers, sequenced tissues, and animal life stages at the time of tissue extraction are provided in Table 1. Our deep sequencing revealed 152,462 transcripts, significantly more than the published counts from other carp genomes, while identifying fewer genes and fewer lncRNAs (6,072) that were longer on average.

**Table 1:** Comparison of genome annotation information. This table includes reference numbers, sequenced tissues, and the life stages of animals at the time of tissue extraction. It lists the number of genes, transcripts, identified lncRNAs, and their median, minimum, and maximum lengths. Our annotation results for crucian carp are indicated in blue.

|  | Genes | Transcripts | lncRNAs | Median length (bp) | Min length (bp) | Max length (bp) | Sequenced tissue | Life stage | NCBI RefSeq assembly |
| --- | --- | --- | --- | --- | --- | --- | --- | --- | --- |
| Crucian carp (*Carassius carassius*) | 45,667 | 152,462 | 6,072 | 1,256 | 198 | 23,112 | Brain | Adult | - |
| Crucian carp (*Carassius carassius*) | 67,308 | 101,854 | 5,793 | 622 | 112 | 5,993 | Multiple* | Juvenile | GCF_963082965.1 |
| Common carp (*Cyprinus carpio*) | 53,084 | 93,194 | 7,684 | 756 | 64 | 15,584 | Muscle | Adult | GCF_018340385.1 |
| Goldfish (*Carassius auratus*) | 75,736 | 123,405 | 8,991 | 795 | 78 | 12,752 | Muscle | Juvenile | GCF_003368295.1 |

\*Annotation was based on varied tissues, different life stages, and RNA sequencing data from *Carassius auratus*.

### Classification of Differentially Expressed LncRNAs and the Nearby Genes

In addition to comparing the length distribution of the DETs with that of all transcripts (Figure 1B), we also compared the length density distributions of the DElncRNAs (1,321) and the identified lncRNAs with sufficient read counts (5,016) (Figure 2A). Both exhibited unimodal, right-skewed shapes with overlapping central tendencies, indicating that the most frequent transcript lengths were identical for both lncRNAs and DElncRNAs. While the DElncRNA density curve showed slightly more variability towards longer transcripts, the overall similarity in their distribution was evident. All lncRNAs started at around 0.2 kb (the cutoff for classification as lncRNA). Of the 1,321 identified DElncRNAs, the median length was 1,672 bp, ranging from 199 bp to 20,462 bp (Figure 2A). Utilizing FEELnc, we successfully predicted interaction partners for 5,951 out of the total 6,072 identified lncRNAs. Among these, 1,306 were DElncRNAs with sufficient read counts. The FEELnc classifier predicted multiple potential interaction partners, with a maximum of 27, a minimum of one, and a median of three partners per DElncRNA, as illustrated in Figure 2B. FEELnc also suggested the best-predicted interaction partner for each lncRNA, allowing us to filter for that and plot the distance between lncRNAs and their predicted interaction partners (Figure 2C). The median distance from lncRNA to nearest genes was 2049 kb. Most DElncRNAs had interaction partners at close distances, up to around 50kb downstream and slightly further upstream. For a detailed classification of lncRNA and DElncRNA subtypes and their genomic locations relative to interaction partners, rows without strand information were removed from the dataset, resulting in 4,216 lncRNAs and 1,129 DElncRNAs retained for analysis. Their distributions were similar, mainly differing in their counts (Figure 2D-H). Approximately 40% of the lncRNAs were located within genes, while around 60% were intergenic (Figure 2D). Most genic lncRNAs were nested within their interaction partner genes, predominantly on the antisense strand. Some were on the sense strand, and some lacked specific strand information. The second most common configuration was overlapping, where the lncRNA partially overlapped with exons or intronic regions of the partner gene. Also, most lncRNAs were on the antisense strand, with fewer on the sense strand. The rarest configuration was containing, where the lncRNA contained the RNA partner transcript. These were mainly on the antisense strand, with a small fraction on the sense strand. Although the nested subtype was predominant among the lncRNAs, the DElncRNAs exhibited a relatively higher proportion of containing and overlapping subtypes (Figure 2E). Most intergenic lncRNAs and DElncRNAs were on the same strand as their RNA interaction partner, indicating transcription in the same orientation (Figure 2F). The least common DElncRNAs exhibited convergent interactions with tail-to-tail orientation, while divergent interactions were characterized by head-to-head transcription with their partners (Figure 2F).

**Figure 2:**
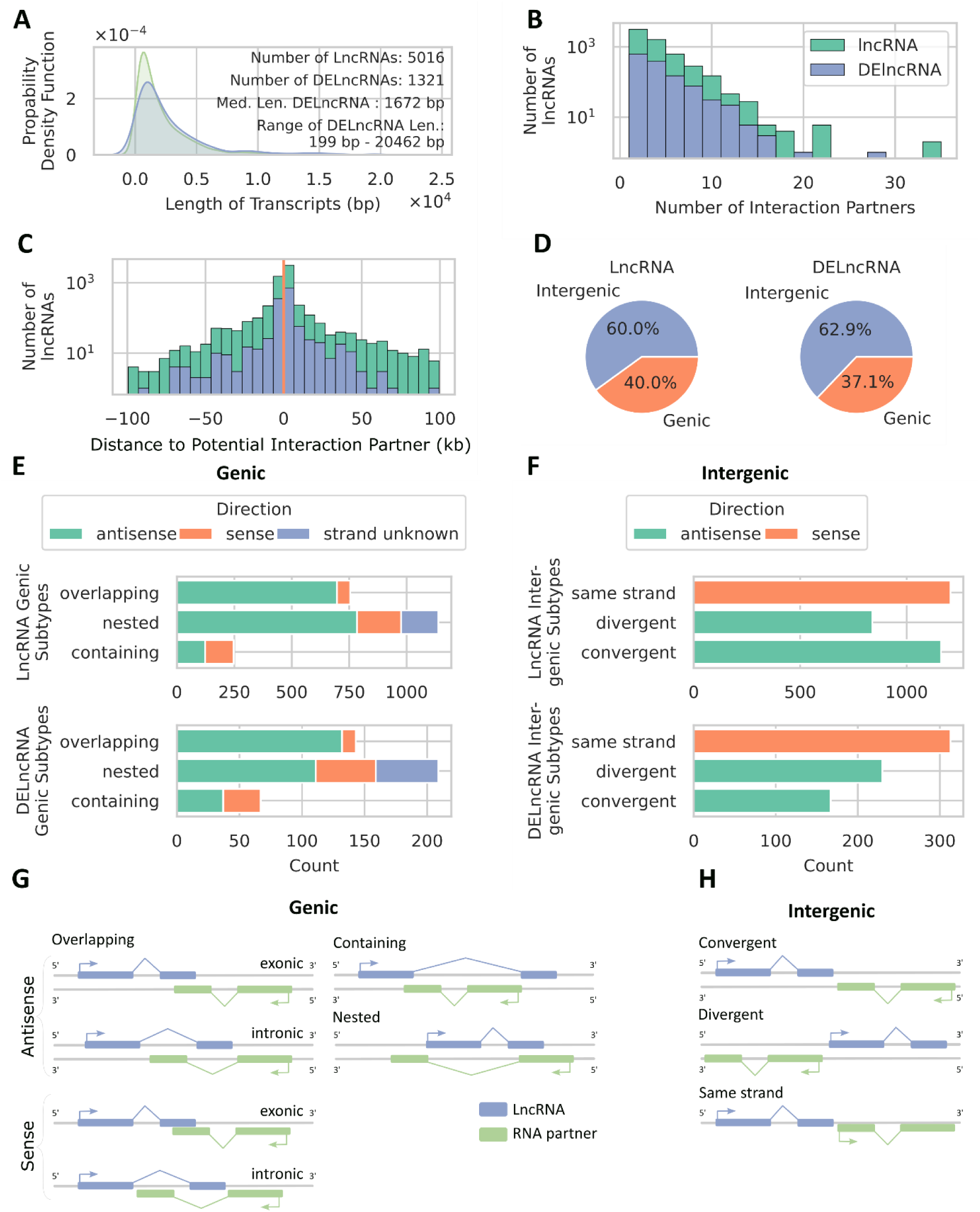
Classification of lncRNAs. A: The density distribution between transcript length and quantity is depicted for lncRNAs (green) and DElncRNAs (blue). Additionally provided is the number of transcripts assessed, from which the density distribution was derived, alongside the median length and range of DElncRNAs. B: Number of Interaction partners for all lncRNAs (green) and DElncRNAs (blue). C: Distance to interaction partners for all lncRNAs (green) and DElncRNAs (blue). Positive values indicate downstream partners, while negative values indicate upstream partners. D: Distribution of lncRNAs in genic and intergenic regions for all lncRNAs and DElncRNAs. E-F: Stacked bar plot of lncRNA subtypes in the genic (E) and intergenic (F) context (see G and H for subtype explanations) for lncRNAs and DElncRNAs. G: Graphical illustration of lncRNA subtypes in the genic context (adapted from Wucher et al. (2017)). Overlapping: the lncRNA partially overlaps the RNA partner transcript; Containing: the lncRNA contains the RNA partner transcript; Nested: the lncRNA is contained in the RNA partner transcript. H: Graphical illustration of lncRNA subtypes in the intergenic context (adapted from Wucher et al. (2017)). Same strand: the lncRNA is transcribed in the same orientation; Divergent: the lncRNA is transcribed in head-to-head orientation; Convergent: the lncRNA is oriented in tail-to-tail orientation.

### Identity and Differential Expression of Potential Interaction Partners

To further elucidate the potential interaction partners of DElncRNA, their mRNA abundance was visualized in a clustered heatmap (batch 1) (Figure 3A), which revealed four distinct clusters. Beginning from the top of the plot, the first cluster showed a decrease in mRNA abundance during normoxia and reoxygenation. Conversely, the second cluster displayed high mRNA abundance in normoxia and low mRNA abundance in anoxia and reoxygenation. The third cluster exhibited low abundance in both normoxia and anoxia, peaking during reoxygenation. The final cluster started with low abundance in normoxia, increased during anoxia, and remained stable during reoxygenation. A PCA was conducted on mRNA abundance data displayed in the heatmap to verify clustering accuracy (since only row clustering was used to create the clustered heatmap), revealing alignment with experimental conditions and no significant sex-specific gene expression differences (Figure 3B). Of the 45,667 protein-coding genes exceeding the ten-read threshold, differential expression was analyzed across three comparisons (N to A, N to R, A to R). Filtering for predicted interaction partners of DElncRNAs subsequently revealed 291 differentially expressed partners (Figure 3C). As observed with lncRNAs, the most substantial changes occurred in the N to A and the N to R comparison and the fewest in A to R. To infer the potential biological functions of differentially expressed DElncRNA predicted interaction partners, a GO analysis was performed across the comparisons: N to A, N to R, and A to R. It revealed that among 291 differentially expressed interaction partners, no GO terms significantly overrepresented after Bonferroni correction. Given the observed expression changes in both DElncRNAs and their predicted interaction partners, we also investigated the potential correlation between their expression levels (Figure 3D), and a positive correlation could be identified (Pearson correlation N to A: 0.35 (p-value: 9.47· 10^-31^), N to R: 0.57 (p-value: 8.87· 10^-87^), A to R: 0.35 (p-value: 1.34· 10^-30^)). This prompted us to generate detailed mRNA abundance plots for the top three most significantly changing DElncRNAs and their predicted interaction partners, as determined by the smallest adjusted p-values and largest log2(fold-changes), without considering their experimental condition (Figure 4, 5). Table 2 details DElncRNA predicted protein-coding interaction partners, including gene ID, protein name, and description. The DElncRNA *STRG8312.1* (Figure 4A) showed increased mRNA abundance during anoxia, returning to normoxic levels upon reoxygenation. Its interaction partner, TM275 (Figure 4B), also had a significant increase in mRNA abundance in anoxia compared to normoxia. *STRG.15579.1* (Figure 4C), exhibited a similar pattern, peaking during anoxia and normalizing in reoxygenation, while its interaction partner, ZN729 (Figure 4D), showed no significant changes compared to normoxia. Lastly, *STRG.9408.4* (Figure 4E), had a very pronounced mRNA abundance increase during anoxia, exceeding a ten-fold change. Although its mRNA abundance levels decreased significantly in reoxygenation, they remained higher than in normoxia. Its interaction partner, SUFU, experienced a significant mRNA abundance decrease during anoxia but returned to normoxic levels in reoxygenation (Figure 4F). The DElncRNA *STRG.9332.1* (Figure 5B) (associated with SRTD2 (Figure 5A)) showed no significant change during anoxia but had a notable increase in mRNA abundance during reoxygenation compared to normoxia. Surprisingly, the mRNA abundance levels of its interaction partner SRTD2 rose considerably in anoxia. The DElncRNA *STRG.27218.4* (Figure 5D) displayed a significant increase in mRNA abundance during anoxia, maintaining elevated levels in reoxygenation. Its interaction partner, MYCB (Figure 5C), also showed increased abundance in anoxia but significantly decreased upon reoxygenation, displaying a divergence in expression patterns compared to its DElncRNA. For DElncRNA *STRG.1182.2* (Figure 5F), the mRNA abundance pattern resembled that of its interaction partner (PMM1; Figure 5E), both showing increased levels during anoxia, followed by a reduction in reoxygenation (Figure 5).

**Figure 3:**
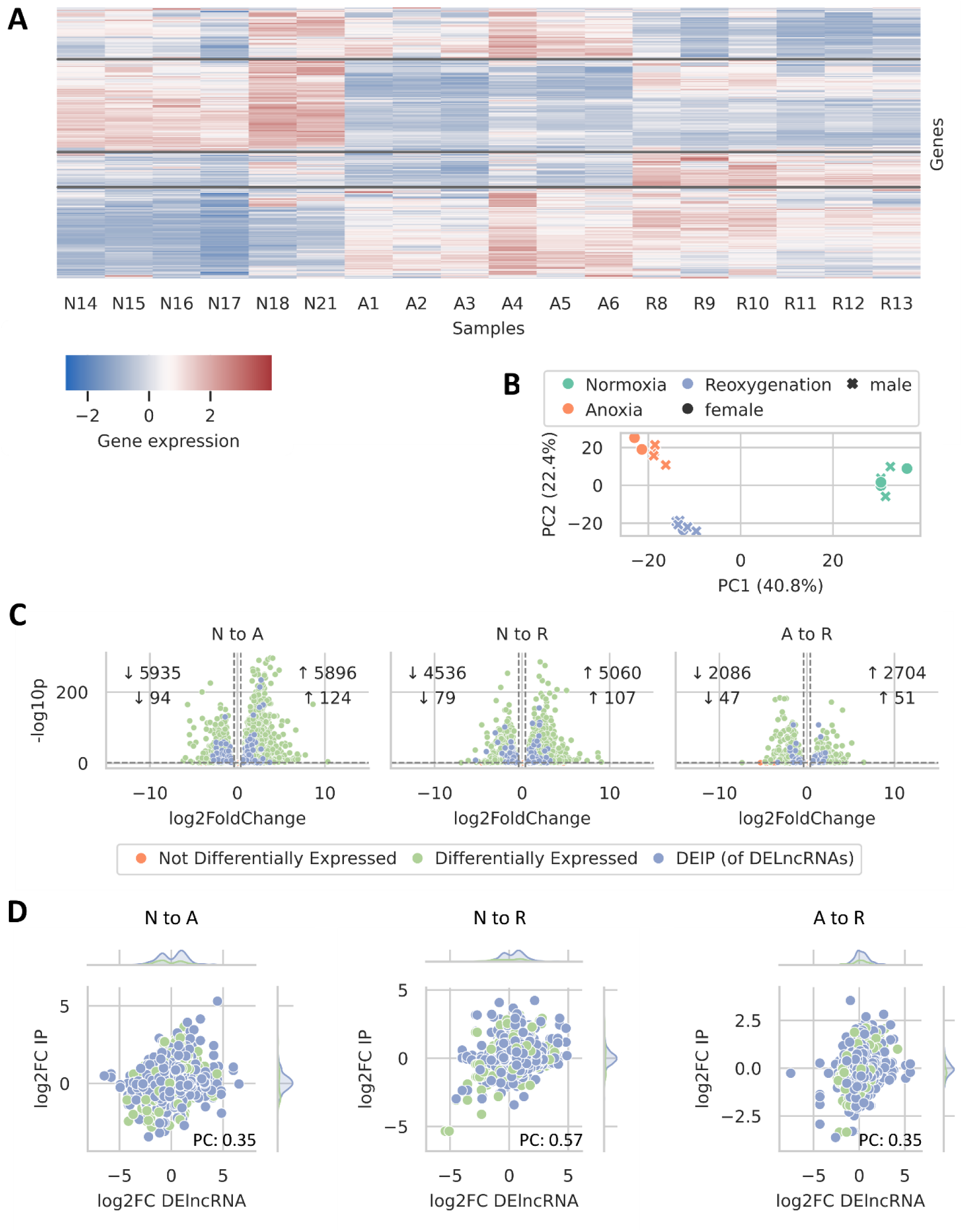
Characterization of Expression Changes in DElncRNA Predicted Interaction Partners. A: Cluster map illustrating the abundance patterns of the DElncRNA predicted interaction partners. Rows normalized with z-score scaling accentuating relative changes, marking gene clusters with similar expression profiles while retaining the column order from normoxia to anoxia and reoxygenation. Sample conditions are provided under normoxia (N), anoxia (A), and 24-hour reoxygenation (R), with corresponding identifiers following each condition. B: PCA revealing sample clustering for the predicted interaction partners of DElncRNA (shown in the cluster map), using six biological replicates across three conditions (2013 samples). C: Volcano plot highlighting significantly differentially expressed predicted interaction partner genes (DEIPs) (of DElncRNA) between treatment conditions, with an adjusted p-value < 0.05 and |log2(fold-change)| ≥ 0.38. The top numbers in each inset denote changes in all differentially expressed genes (green), while the bottom numbers indicate DEIPs (blue) with increased (↑) or decreased (↓) expression. D: Joint plot conveying the correlation between abundance changes (log2 fold changes) in DElncRNAs and their predicted interaction partners (IP), with histograms on the top and right margins illustrating sample abundance in the corresponding plot areas. DElncRNAs with non-differentially expressed partners (green) and differentially expressed partners (blue). PC: Pearson correlation.

**Figure 4:**
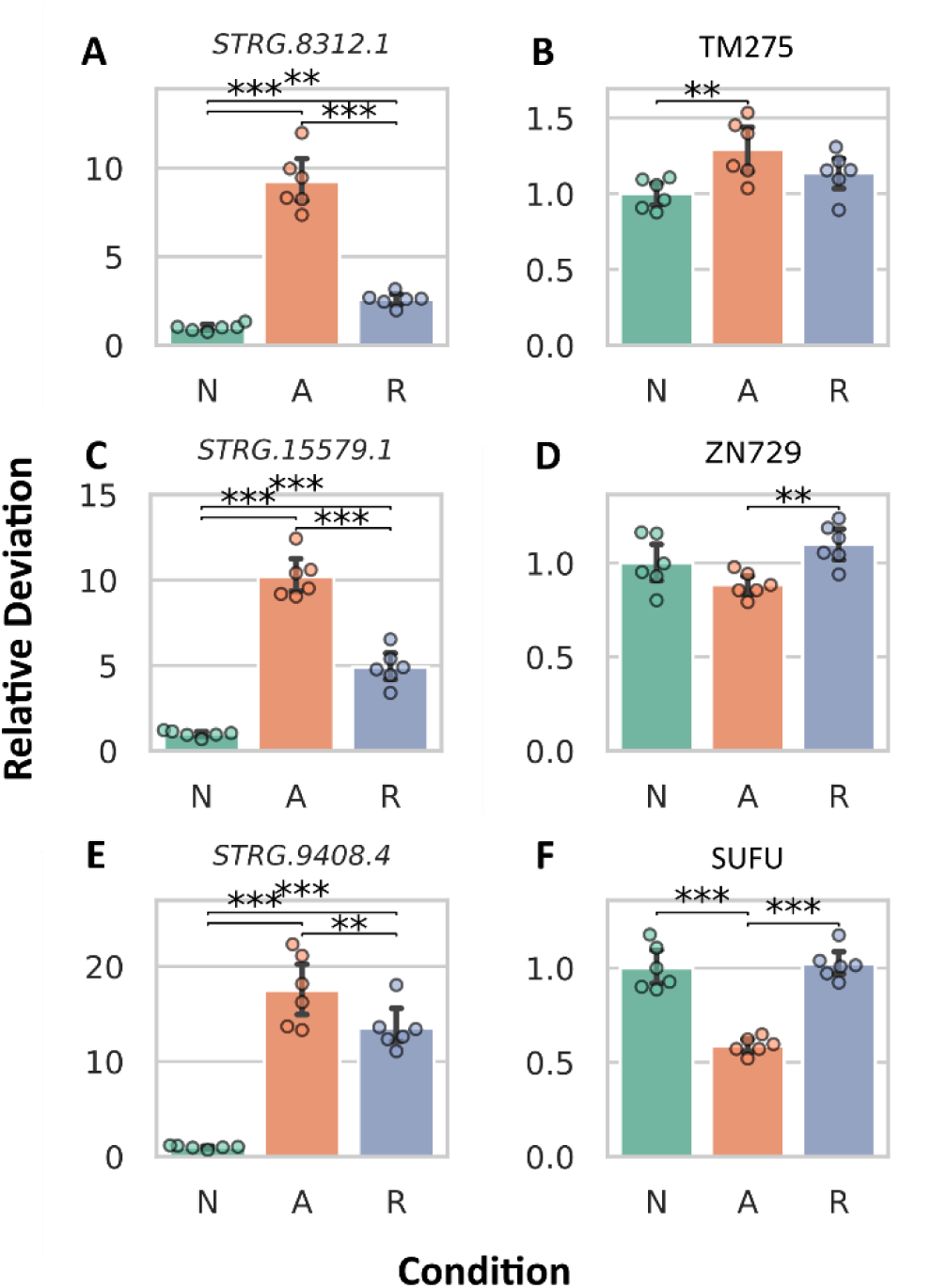
Abundance changes in most differentially expressed DElncRNAs (left column) and their predicted interaction partners (right column). They were selected based on the lowest adjusted p-values and highest log2(fold-changes). The top three candidates with adequate read counts (exceeding 50 across all samples) are depicted.

**Figure 5:**
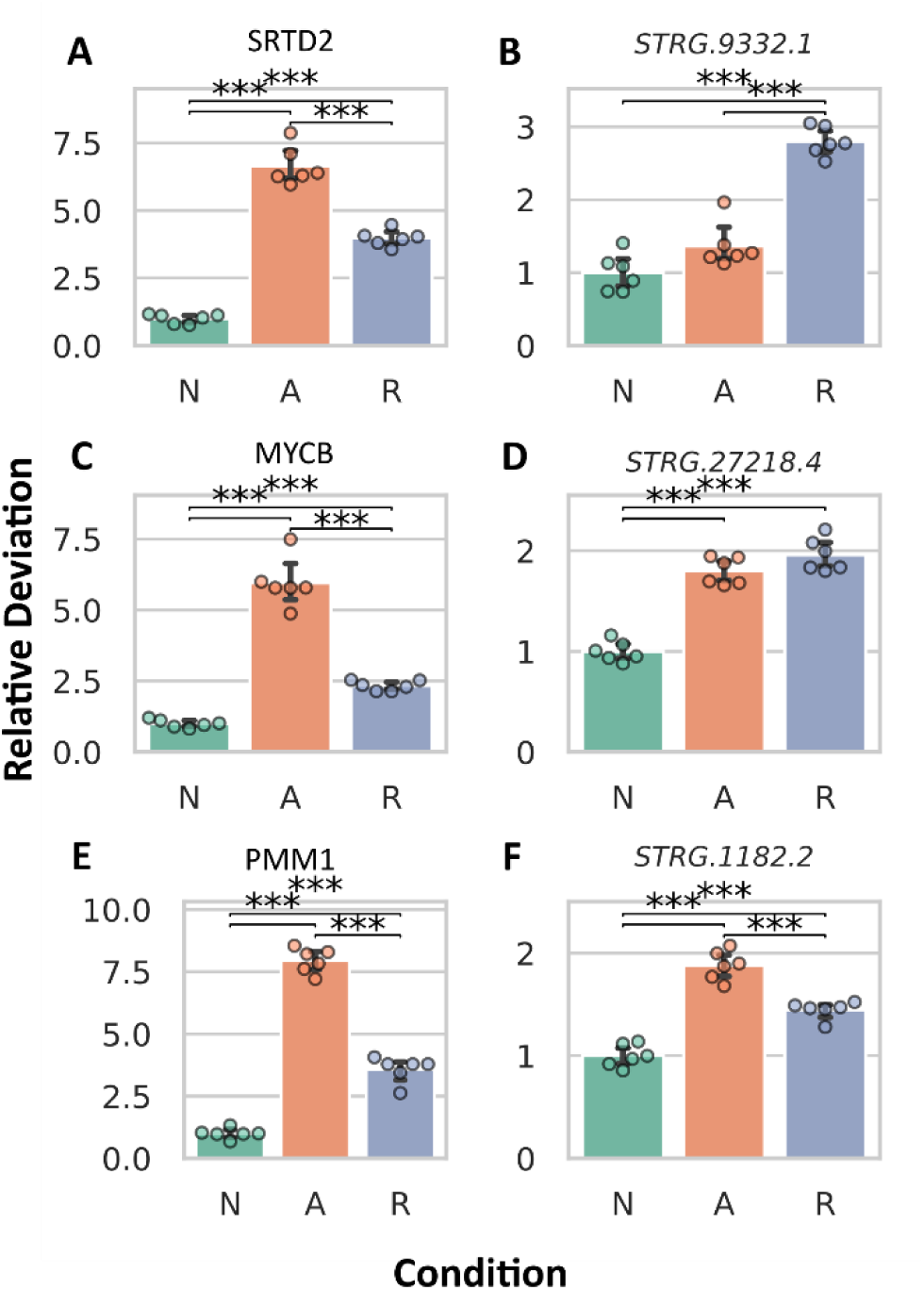
Abundance changes in the most differentially expressed predicted interaction partners (left column) and their DElncRNAs (right column). They were selected based on the lowest adjusted p-values and highest log2(fold-changes). The top three candidates with adequate read counts (exceeding 50 across all samples) are depicted.

**Table 2:** Identified homolog proteins in the SwissProt database of the most differentially expressed DElncRNA predicted interaction partner genes.

| Gene ID | Symbol | Description |
| --- | --- | --- |
| ccar_ua10-g9980 | TM275_MOUSE | Transmembrane protein 275 |
| ccar_ua20-g18868 | ZN729_HUMAN | Zinc finger protein 729 |
| ccar_ua11-g11305 | SUFU_HUMAN | Suppressor of fused homolog (SUFUH) |
| ccar_ua11-g11216 | SRTD2_HUMAN | SERTA domain-containing protein 2 (Transcriptional regulator interacting with the PHD-bromodomain 2) (TRIP-Br2) |
| ccar_ub10-g32843 | MYCB_DANRE | Transcriptional regulator Myc-B (c-Myc-B) |
| ccar_ua01-g1406 | PMM1_MOUSE | Phosphomannomutase 1 (PMM 1) (EC 5.4.2.8) |

## Discussion

LncRNAs are emerging as pivotal regulators of various cellular mechanisms, including gene expression (Du, Zhang, Wu, You, & Dong, 2023). This study investigated whether the abundance of lncRNA transcripts is affected by anoxia, and thus their potential involvement in regulating transcriptional responses to anoxia in the brain of the crucian carp, an anoxia-tolerant species with previously uncharacterized lncRNAs. We identified the lncRNAs using CPC2, CPAT, CNCI, and FEELnc, resulting in 6,072 lncRNAs (Figure 1C). Multiple programs were chosen to leverage diverse algorithms (for details, see Methodological Considerations below). We observe that the individual programs identified lncRNAs within a similar magnitude in numbers, reinforcing the reliability of our results. While the numbers do not completely align, this is expected due to the use of both alignment-based and alignment-free algorithms (for details, see Methodological Considerations). Comparing our transcriptome (this study) and genome (Valencia-Pesqueira, Hoff, Tørresen, Jentoft, & Lefevre, 2025) By comparing our assembly results with those of other published genomes and annotations, we identified fewer genes but more transcripts using our Stringtie approach. This likely results from our deep sequencing, which recovered less abundant transcripts. We identified 152,462 transcripts, significantly more than the 101,854 transcripts from 67,308 genes found in the published (farmed) crucian carp genome (GCF_963082965.1). For the common carp (GCF_018340385.1), 93,194 transcripts were identified from 53,084 genes, and in the goldfish (GCF_003368295.1), 123,405 transcripts were found from 75,736 genes. We identified a smaller number of lncRNAs, with 6,072, compared to approximately 5,000 to 9,000 in the other assemblies. Our identified lncRNAs had a median length of 1,256 bp, significantly longer than the 622 - 795 bp range in other assemblies. Our maximum lncRNA length was 23,112 bp, compared to 5,993 bp in farmed crucian carp, 15,584 bp in common carp, and 12,752 bp in goldfish. Notably, the assemblies also noted lncRNAs shorter than 200 bp, likely due to classification at the transcript level, resulting in transcript lengths under 200 bp. It is essential to consider that sequencing from different tissues and life stages, as well as differences in the genome annotation approach, may also account for some of the observed variation in numbers. We utilized StringTie for transcriptome assembly, unlike the published genomes, which have been annotated using other methodologies (i.e. NCBI Eukaryotic Genome Annotation Pipeline).

Having established a robust identification of lncRNAs, we further investigated the differential expression of lncRNAs at the transcript level (Figure 1D). At the transcript level, most changes in abundance occurred upon entering anoxia, with a downregulation of 492 transcripts. These lncRNAs might be involved in energy preservation through reduction of transcription, aligning with the known metabolic downregulation in crucian carp during anoxia (Dahl, Johansen, Nilsson, & Lefevre, 2021). Additionally, 491 transcripts were upregulated in anoxia, suggesting their potential role in regulating anoxia-specific genes. The fewest DElncRNAs were observed during the transition from anoxia to reoxygenation, implying less involvement of lncRNA in the response to reoxygenation, or potentially slower turnover of lncRNAs compared to protein-coding transcripts, which exhibited a pronounced response from anoxia to reoxygenation (Figure 1D). It is also possible that the crucian carp maintains anoxic adjustments for a while in anticipation of further anoxic episodes. When comparing N to R, downregulated transcripts were halved compared to N to A, while upregulated transcripts increased to 513, surpassing the N to A comparison. Another lncRNA study investigated gonad development in common carp, identifying 106,932 mRNA transcripts, including 14,199 lncRNAs, with 124 (0.87%) differentially expressed (Song, Wang, Zhu, & Dong, 2019). In contrast, our study identified a higher percentage of DElncRNAs, ranging from 1.30 - 3.14% on the transcript level in all comparisons. Song et al. (2019) solely utilized CPC for lncRNA identification with a different cut-off for the classification of lncRNAs by the algorithm (>0) and used edgeR for DEG analysis (|log2(fold-change)| >1 and p-value <0.05), which differed from our methods (Song, Wang, Zhu, & Dong, 2019). Song et al. (2019) found more transcripts and lncRNAs than the published NCBI genome of common carp (GCF_018340385.1), likely due to working on embryos and whole organisms, whereas the published annotation used juvenile muscle tissue (Song, Wang, Zhu, & Dong, 2019). Since the abundance of lncRNAs can be highly tissue-specific, this could explain their higher lncRNA count. When examining the DElncRNA transcripts (Figure 1E), we observed that out of the 1,321 transcripts, 348 (26.34%) were solely found in the N to A comparison, suggesting that those transcripts had returned to normoxic levels after just 24 hours of reoxygenation. In the N to A and N to R comparison, 417 (31.57%) were differentially expressed in both, indicating that these DElncRNAs were differentially expressed in the N to A comparison and remained so after 24 hours of reoxygenation. Further, in anoxia, 176 (13.32%) DElncRNAs could be essential for maintaining the necessary activity of anoxia-related genes. The 70 DElncRNAs (5.30%) identified solely in the A to R comparison could likely play a role in transitioning from anoxia to reoxygenation, potentially reactivating normoxia genes and deactivating anoxia genes. The 42 delncRNAs (3.18%) common across all comparisons indicate more complex changes in abundance during anoxia and reoxygenation. Upon reoxygenation, their abundance changes or returns towards normoxic levels, yet they remain significantly differentially expressed.

When comparing all lncRNAs with the differentially expressed ones (Figure 2A), we observed largely similar patterns in length distribution, numbers of predicted potential interaction partners distance (Figure 2B-C), genic and intergenic occurrence (Figure 2D), and subtype classification (Figure 2E-G). This indicates that the DElncRNAs do not appear to have verz different characteristics compared to lncRNAs in general, and hence, we will discuss lncRNA characteristics collectively without distinguishing whether the lncRNAs were differentially expressed or not. We found that for a single DELncRNA, the classifier module of the FEELnc predicted multiple potential interaction partners, with a maximum of 27 (Figure 2B). This is consistent with the literature, which states that one lncRNA can regulate multiple target genes, and multiple lncRNAs can collaborate to regulate a single gene (Park & Kim, 2023). However, most lncRNAs had only one potential interaction partner. In terms of proximity, many of our identified lncRNAs had interaction partners nearby, while fewer lncRNAs exhibited interactions over greater distances. Luo et al. (2019) investigated lncRNA expression in differently colored skins of koi carp (*Cyprinus carpio L.*) - black, white, and red - identifying 77,907 lncRNAs, of which 92 were DElncRNAs. They identified putative lncRNAs using NONCODE, filtering for transcripts >200 nucleotides and removing those with ORFs >300 bp. The remaining transcripts were aligned to the NCBI database to exclude known proteins, and CPC was used to classify transcripts with scores <0 as novel lncRNAs. *Cis* interactions involve lncRNAs regulating neighboring target genes within 10 - 100 kb upstream or downstream (Luo, et al., 2019). They found that 70 lncRNAs acted on 107 target mRNAs in *cis* (= lncRNAs regulating genes on their originating chromosome), while 79 lncRNAs acted on 41,625 targets in *trans*, demonstrating that a single lncRNA can connect to numerous mRNAs. To predict potential interaction partners, they searched for coding genes within 10-100 kb of lncRNAs (*cis*-acting) and calculated Pearson correlation coefficients (p < 0.05, R ≥ 0.95) between lncRNAs and coding genes (*trans*-acting; the method used to identify these targets was not specified) using a custom script. In contrast to *cis*, *trans* interactions occur over considerable distances, spanning megabases or involving entirely different chromosomes. Due to the composition of the FEELnc algorithm, *trans*-acting lncRNAs were not assessed directly in this study, as the algorithm does not classify them. But considering FEELnc only identifies the nearest transcripts from the reference annotation and the distances observed in this study are all below 100 kb (maximum: 99,979 bp), it suggests that the vast majority of identified lncRNAs in crucian carp are *cis*-acting, even though the existence of *trans* interactions cannot be excluded. Additionally, *cis*-acting lncRNAs are generally more common due to their localized functionality, reducing the likelihood of dilution through diffusion or transport to other cellular compartments (Kazimierczyk & Wrzesinski, 2021; Park & Kim, 2023; Muret, et al., 2017). The proportion of lncRNAs in the intergenic context was 1.5 times higher than in the genic context (Figure 2D). This is plausible because the entire genome comprises more intergenic than genic sequences. In the intergenic context, most lncRNAs were located on the same strand as their potential interaction partner, while convergent and divergent lncRNAs were less common. In the genic context, overlapping and nested lncRNAs were mainly on the antisense strand, with some on the sense strand, and many nested lncRNAs lacked strand information. Containing DElncRNAs were primarily on the antisense strand, with a few on the sense strand (Figure 2E-H). Given the limited knowledge about lncRNAs, we have no basis for comparison. We cannot determine if the identified subclasses and their distribution are typical for a specific group of animals (such as cyprinids, teleost fish, or vertebrates).

To assess whether changes in the abundance of DElncRNAs were also reflected in the abundance of their predicted interaction partners, we constructed a clustered heatmap (Figure 3A), revealing multiple clusters with distinct abundance shifts under different oxygen availability conditions. Subsequently, a differential expression analysis quantified the extent of abundance variation, identifying numerous differentially expressed interaction partners (Figure 3C). We performed a GO analysis to gain insight into the functional roles of DElncRNA interaction partners that were also differentially expressed. The analysis revealed that among the differentially expressed interaction partners, no GO terms were significantly overrepresented after Bonferroni correction of the p-values. To explore whether increases in DElncRNA expression correlated with the expression of their predicted interaction partners, we plotted their log2(fold-changes) (Figure 3D) against one another, and found positive Pearson correlation coefficients (N to A: 0.35, N to R: 0.57, A to R: 0.35), indicating positive correlations and thus potential co-expression of DElncRNAs with the closest nearby gene, or possibly regulation of the mRNA abundance of the nearby gene. Furthermore, we highlighted the three most differentially expressed DElncRNAs, identified by the lowest adjusted p-values and largest log2(fold-changes), along with their interaction partners (Figure 4 - 5). Our analysis revealed that the three most DElncRNAs had significantly higher mRNA abundance in anoxia compared to normoxia, with levels remaining elevated even during reoxygenation. The mRNA abundance of their predicted interaction partners varied: The mRNA predicted to code for transmembrane protein 275 (TM275) increased, the mRNA predicted to code for zinc finger protein 729 (ZN729) remained unchanged, and the mRNA predicted to code for suppressor of fused homolog (SUFU) exhibited an inverse correlation in anoxia. TM275 [NCBI Gene ID: 76237] is predicted to reside in the membrane, suggesting its role in nutrient import and waste export, which are essential for cell survival. ZN729 [NCBI Gene ID: 100287226] is predicted to facilitate DNA and zinc ion binding, and may play a role in regulating transcription by RNA polymerase II. SUFU [NCBI Gene ID: 51684] negatively regulates the Hedgehog signaling pathway, which is important for growth, development, and stress responses. In anoxia, this could aid cellular survival and adaptive responses by managing metabolic processes. One could speculate that *STRG8312.1* may enhance TM275 expression, while *STRG.15579.1* might exert no influence on ZN729, or ZN729 could be suppressed by other factors. Additionally, *STRG.9408.4* and SUFU may demonstrate the potential for lncRNAs to function as repressors of mRNA abundance. However, without a comprehensive understanding of these relationships, it is difficult to ascertain whether they represent genuine correlations or are part of a more complex network of interactions that remains to be uncovered. Therefore, interpreting these findings poses challenges, as the possibility of false correlations cannot be dismissed.

The analysis of lncRNAs and their most differentially expressed interaction partners revealed that *STRG.9332.1* showed no increase in mRNA abundance in anoxia, while its interaction partner, the SERTA domain-containing protein 2 (SRTD2), exhibited a significant increase. SRTD2, known for regulating fat storage by downregulating genes involved in adipocyte lipolysis, thermogenesis, and oxidative metabolism (Hsu, 2001), is of particular interest given the reliance of crucian carp on glycogen reserves during oxygen deprivation. In contrast, the other two DElncRNAs (*STRG.27218.4, STRG.1182.2*) showed increased mRNA abundances in anoxia, which was mirrored by their partners, transcriptional regulator Myc-B (MYCB) and phosphomannomutase 1 (PMM1). MYCB [NCBI Gene ID: 393141] functions as a DNA-binding transcription factor specific to RNA polymerase II, playing a role in transcription regulation. PMM1 [NCBI Gene ID: 5372] catalyzes the conversion of D-mannose 6-phosphate to D-mannose 1-phosphate, a precursor for GDP-mannose, which is necessary for N-linked glycosylation and the production of glycoproteins. Notably, for *STRG.27218.4*, its mRNA abundance remained unchanged during reoxygenation, while the mRNA abundance of MYCB significantly decreased. Only *STRG.1182.2* and its partner PMM1 displayed a consistent pattern in abundance changes. However, as mentioned above, we acknowledge the potential for false correlations, as the complexity of the regulatory network and other influencing factors remain poorly understood.

This detailed characterization of lncRNAs in the brain of crucian carp was conducted in the context of anoxia exposure, focusing on the DElncRNAs, constituting approximately 21.76% of the identified lncRNAs. The activity and regulatory roles of the remaining lncRNAs remain unknown. Literature suggests that lncRNAs are active during development and involved in responding to various stressors and diseases (Bridges, Daulagala, & Kourtidis, 2021). Thus, while the analyzed subset may play a role in regulating the anoxia response, the other lncRNAs could play roles in developmental processes, responses to other stressors, or the development and progression of diseases. It should be noted that teasing apart cause and effect regarding the similarity in responses of lncRNAs and their interaction partners (i.e. the positive correlation) is challenging. Without further mechanistic studies, it cannot be ruled out that both the lncRNA and their partner are affected by upstream factors, leading to co-expression, especially when the lncRNA and the partner are located very close together and on the same strand. The end result is, of course, the same – altered abundance of the interaction partner - and co-expression rather than direct regulation is an equally intriguing possibility, as it might indicate a role for the lncRNA in stabilization or translation of the partner gene.

Our study aligns with the emerging role of lncRNAs as crucial regulators in various cellular mechanisms, including gene expression regulation. We employed multiple algorithms to identify 6,072 lncRNAs that were previously unknown in crucian carp. Differential expression analysis revealed significant transcriptional changes of lncRNAs during anoxia, underscoring the potential involvement of lncRNAs in transcriptional adjustments during the transition into and recovery from anoxia. Although the roles of many lncRNAs remain unknown, the activity of their predicted interaction partners under anoxia highlights their possible regulatory significance. This study enhances our understanding of lncRNAs in crucian carp and their broader roles in cellular regulation under stress conditions.

## Methodological Considerations

Our analysis utilized poly(A) tail capturing to enrich polyadenylated RNAs, a common method for targeting mRNA over the much more abundant ribosomal RNA. This method employs poly(T) oligos to selectively extract RNAs with poly(A) tails, enhancing specificity and reducing RNA pool complexity by focusing on mRNAs and polyadenylated lncRNAs. While this technique improves the identification of polyadenylated lncRNAs, it inherently biases the analysis towards these transcripts, potentially missing non-polyadenylated lncRNAs. This bias is important to consider when interpreting and utilizing the data and information gathered in this study (Mattick, et al., 2023). LncRNA identification using CPC2, CPAT, CNCI, and FEELnc resulted in the detection of 6,072 lncRNAs (Figure 1C). We selected multiple programs to leverage diverse algorithms. Given that lineage-specific and less conserved lncRNAs in non-model organisms are more challenging to identify, we used species-neutral, alignment-free tools, as alignment-based methods can be biased towards conserved protein-coding genes and struggle with species-specific lncRNAs (Wang, et al., 2013). Alignment-based methods are effective for conserved protein-coding genes but can struggle with novel transcripts, especially lineage-specific lncRNAs. This can result in misclassifying unknown protein-coding genes of the investigated species as non-coding due to insufficient similarity to conserved genes. For instance, only 29 out of 550 identified lncRNAs in zebrafish showed detectable sequence similarity with mammalian orthologs (Wang, et al., 2013). CPAT, CNCI, and FEELnc are alignment-free methods, while the alignment-based CPC2 was included due to its newer design and species-neutral capabilities, making it suitable for non-model organisms like crucian carp. A disadvantage of CPAT is its reliance on zebrafish data for model training rather than solely on intrinsic sequence composition features (Wang, et al., 2013). CNCI uses a general vertebrate model, and FEELnc employs a Random Forest model trained on features like multi k-mer frequencies and relaxed open reading frames. We used these algorithms together to balance the strengths and limitations of several algorithms. A similar strategy was employed by Weikard et al. (2018), who investigated lncRNAs in the jejunum (part of the small intestine) transcriptome of German Holstein calves fed two different milk diets. Using four programs - FEELnc, CNCI, PLEK, and PLAR - they identified 1,812 lncRNAs, of which 9 were DElncRNAs. Like our study, their tools were divided into alignment-free (CNCI, PLEK, and FEELnc) and alignment-dependent (PLAR) algorithms. However, we did not use PLEK and PLAR in our study due to their predisposition for mammalian organisms (Weikard, et al., 2018).

## Author contribution

**Magdalena Winklhofer**: Formal analysis, Methodology, Investigation, Data curation, Visualization, Writing-Original Draft, Writing-Review and Editing; **Göran E. Nilsson**: Funding acquisition (supporting), Resources, Writing-Review and Editing; **Sjannie Lefevre**: Conceptualization, Methodology, Resources, Writing-Review and Editing (lead), Supervision, Project administration, Funding acquisition (lead).

## Data availability and resources

All raw RNA sequencing data from batch 1 dataset are deposited in the NCBI Sequence Read Archive (SRA) under BioProject ID PRJNA386629. The batch 2 dataset is deposited in the NCBI SRA under BioProject Accession PRJNA1163668. The genome sequence and annotation data were obtained from DataverseNO (https://doi.org/10.18710/GXMSUH). Supplementary Material for the present study has been deposited in DataverseNO: https://doi.org/10.18710/1EIOVF, and scripts are available in the GitHub repository LncRNA (https://github.com/MagdalenaWinklhofer/paper_lncRNA.git).

## Acknowledgments

The authors express their gratitude to the staff at the Norwegian Sequencing Center for carrying out the RNA library preparation as well as the RNA sequencing. They also thank Sigma2 - Norway’s National Infrastructure for High-Performance Computing and Data Storage - for facilitating data analysis and the staff at the InVivo facility at the Department of Biosciences for their support in crucian carp husbandry. Additionally, the authors appreciate the assistance of former Lefevre-Nilsson research group member May Kristin Torp with fish collection, anoxia exposure, and sampling, and Cathrine E. Fagernes for assistance with RNA extraction. Special thanks are also extended to the current members of the Lefevre-Nilsson research group (Elie Farhat, Lucie Gerber, Laura Marian Valencia-Pesqueira, and Jenny Lundeberg) for their insightful discussions.

